# PLFest: A Multi-Site Validation of an Open Platform for Visual and Cognitive Assessment

**DOI:** 10.64898/2026.06.22.733892

**Authors:** Boris Penaloza, Marcello Maniglia, Jaap Munneke, C. Shawn Green, Aaron Seitz

**Affiliations:** Department of Psychology, Northeastern University, Boston, MA, USA; Department of Psychology, Rochester Institute of Technology, NY, USA; Department of Psychology, University of Wisconsin Madison, Madison, WI, USA

## Abstract

**Purpose:** To evaluate the feasibility, validity, and scalability of PLFest, an open-source, Unity-based, cross-platform application designed for standardized, multi-site visual and cognitive assessment and training.

**Methods:** Two hundred sixty participants (mean age = 23 years) were recruited across four university sites in the United States. Participants completed a battery of five visual assessments administered through PLFest, including visual acuity, contrast sensitivity, spatial frequency cutoff, contrast sensitivity at spatial-frequency cutoff, and visual search. Five cognitive assessments measuring visuospatial working memory, verbal working memory, fluid reasoning, inhibitory control, and selective attention were also administered. Descriptive statistics and performance distributions were examined and compared with normative data.

**Results:** Visual acuity and contrast sensitivity measures closely matched previously reported normative values obtained using established clinical and psychophysical methods. Spatial frequency cutoff and visual search tasks produced stable threshold estimates while showing substantial inter-individual variability. Performance across all cognitive assessments was consistent with published validation studies of the corresponding tasks. Across the full battery, adaptive procedures demonstrated reliable convergence and generated well-distributed performance measures without evidence of substantial floor or ceiling effects. Importantly, these findings were observed across four geographically distributed testing sites using standardized consumer-grade tablet hardware.

**Conclusions:** PLFest provides reliable and scalable assessment of visual and cognitive function using portable consumer devices. The platform supports standardized data collection across distributed research settings while maintaining performance characteristics consistent with established laboratory and clinical benchmarks. These findings support the use of PLFest as a reliable framework for large-scale studies of vision and cognition.

**Translational Relevance:** By reducing dependence on specialized laboratory infrastructure and trained personnel, PLFest may facilitate broader access to visual and cognitive assessment, enabling large-scale research, screening, and future rehabilitation applications.

## 1. Introduction

Perceptual and cognitive abilities are fundamental to everyday functioning, supporting activities ranging from reading and navigation to decision-making in complex environments. Traditionally, assessment and training of these abilities have relied on specialized laboratory equipment, trained personnel, and highly controlled testing environments. While such approaches provide the precision necessary for psychophysical measurement, they can limit scalability, accessibility, and ecological validity.

Over the past decade, the field of vision science, and in particular perceptual learning––the neural plasticity-based improvement in sensory functions following repeated practice (Sagi, 2011)––, have attracted increasing interest as tools for studying neural plasticity and improving visual function. Laboratory-based perceptual interventions have shown promise in healthy observers and in clinical populations with conditions such as myopia (Tan & Fong, 2008), presbyopia (Polat et al., 2012), amblyopia (Levi & Li, 2009), and central vision loss (Chung, 2011; Maniglia et al., 2016; Maniglia et al., 2020). Despite promising results, progress towards effective training paradigms has been constrained by challenges related to reproducibility and accessibility. Similar to broader concerns within psychology and neuroscience (Ioannidis, 2005), perceptual learning studies often report inconsistent effects across laboratories, with difficulties replicating outcomes across independent research groups (Aberg & Herzog, 2010; Hung & Seitz, 2014; Liang et al., 2015; Xiao et al., 2008; Zhang & Yang, 2014; Zhang & Yu, 2016). Variability in hardware, stimulus presentation, calibration procedures, instructions and experimental implementation likely contributes to these discrepancies. At the same time, the dependence on specialized laboratory infrastructure restricts participation and limits the feasibility of large-scale, multi-site investigations.

Recent advances in consumer-grade tablet technology offer a potential solution to these challenges. Modern tablets provide high-resolution displays, precise timing capabilities, responsive touchscreen interfaces, and sufficient computational power to support sophisticated psychophysical testing (Calabrèse et al., 2018; Rono et al., 2019). Importantly, these devices enable standardized assessment outside traditional laboratory environments while substantially reducing logistical and financial barriers. Digital assessment platforms also offer advantages for clinical and population-based research, including automated administration, scoring, and data management. Such capabilities may facilitate broader access to vision and cognitive assessment while supporting large-scale longitudinal studies and remote data collection. Indeed, recent studies have demonstrated that smartphone-and tablet-based measures of visual function can achieve performance comparable to gold standard clinical assessments, including ETDRS visual acuity and Pelli-Robson contrast sensitivity charts (Bastawrous et al., 2015; Habtamu et al., 2019; Kollbaum et al., 2014).

To capitalize on these technological developments, we recently developed PLFest (Jayakumar et al., 2024), a Unity-based, cross-platform application designed to deliver perceptual learning paradigms and visual assessments across computers, tablets, and mobile devices. PLFest was developed within an open-science framework specifically intended to improve reproducibility, accessibility, and cross-site standardization. By providing consistent stimulus presentation, adaptive testing procedures, automated data handling, and compatibility across widely available hardware, the platform minimizes methodological variability that has historically complicated comparisons across studies and sites.

Beyond improving reproducibility, digital assessment platforms create opportunities to investigate the relationship between perceptual and cognitive function at a scale that has not previously been feasible. Although perceptual and cognitive assessment traditions have often evolved independently, growing evidence suggests that perceptual learning interacts with broader cognitive processes, including attention, working memory, and executive control. Understanding these interactions requires integrated assessment frameworks capable of measuring both perceptual and cognitive performance within a unified testing environment.

In the present study, we extend the validation of PLFest by reporting baseline data from an ongoing multi-site clinical trial involving 260 participants recruited across four geographically distributed university sites. Participants completed a ten-task assessment battery consisting of five visual tasks administered through PLFest, including measures of visual acuity, contrast sensitivity, spatial frequency resolution, and visual search, and five cognitive tasks delivered through the companion Recollect Assessment Battery (Pahor, Seitz, et al., 2022), assessing working memory, fluid intelligence, inhibitory control, and selective attention. Together, these platforms provide a comprehensive framework for integrated perceptual and cognitive assessment.

Our primary objective was to evaluate the feasibility, reliability, and scalability of these tablet-based assessment tools across multiple testing sites. We hypothesized that both platforms would produce stable, well-distributed performance measures consistent with normative benchmarks reported in the literature, despite variability in testing environments. Demonstrating robust performance across geographically distributed settings would provide critical evidence for the external validity of digital psychophysical assessment and support the broader use of portable, low-cost technologies for research, clinical screening, and population health monitoring. Ultimately, such tools may help overcome longstanding barriers to access, enabling more equitable and scalable approaches to vision and cognitive assessment.

## 2. Methods

### 2.1. Participants

A total of 260 observers participated in the study (153 M; mean age = 23 years [SD = 3 years]). The study was conducted across four sites: Northeastern University (NU, N = 73), University of California at Riverside (UCR, N = 173), University of Wisconsin Madison (UW, N = 39), and Mills College at Northeastern University (Mills, N = 11). All participants reported normal or corrected to normal vision. Written informed consent was obtained from all participants, and the study was approved by the IRB of all participating universities.

### 2.2. Exclusion criteria

Participants with anomalous performance thresholds, which likely reflect attentional lapses or failure to understand task instructions, were excluded from analysis. For visual tasks, thresholds were calculated as the average of the last 10 staircase values, and outliers were identified using a two-step procedure combining task-specific cutoffs and removal of values beyond ±3 SD of the group mean. For cognitive tasks, outliers were defined based on implausibly fast reaction times (<200 ms) and scores exceeding 3 SD from the group mean.

### 2.3. Experimental setup

Participants were tested using Apple iPad Pro tablets 2021 with a screen resolution of 2048×2732 pixels and a refresh rate of 120 Hz. Devices were mounted on stands and placed at a viewing distance of 50 cm from the participant. Head and chin rests were used to minimize head movements across participants. Testing was conducted under low-lighting ambient conditions (0.9 cd/m^2^). The iPad background luminance was matched across sites at 75 cd/m^2^. Responses were recorded via a small numpad keyboard attached to the iPad.

Visual assessments were administered through the PLFest App (Jayakumar et al., 2024), a cross-platform UNITY-based application designed to support accessible, reproducible, and open perceptual learning and assessment research. PLFest is compatible with iOS, Android, macOS, and Windows, and supports data collection on tablets, computers, and smartphones, facilitating large-scale, multi-site studies. Cognitive assessments were administered through the Recollect App, a dedicated platform for standardized delivery of cognitive tasks (Pahor, Seitz, et al., 2022).

### 2.4. General procedure

Participants completed a single session lasting approximately one hour. The session included 5 visual psychophysics tasks and 5 cognitive assessments, administered in counterbalanced order across participants and sites. Prior to each task, participants completed a practice phase to familiarize themselves with the stimuli and response requirements. Two-minute breaks were interspersed throughout the various assessments.

### 2.5. Visual assessments tasks

#### 2.5.1. Contrast Sensitivity Function (CSF)

Contrast sensitivity was assessed using centrally presented Gabor patches with a spatial frequency of 6 cycles per degree (cpd), tilted at varying orientations. The 6 cycles/degree stimulus was chosen because it lies near the peak of the human spatial contrast sensitivity function, which generally occurs within the mid-spatial frequency range (approximately 3–10 cycles/degree) under photopic conditions (Campbell & Robson, 1968). Participants indicated the direction of tilt by pressing the corresponding icon in the lower left or right corner of the tablet display (Figure 1A). This task employed a blockwise staircase procedure in which 20 mini-blocks of 6 trials were presented, with each trial corresponding to one of six Gabor orientations (22.5°, 45°, 67.5°, 112.5°, 135°, and 157.5°) in random order. Stimulus contrast decreased (i.e., difficulty increased) when 1 or fewer errors occurred within a block, remained unchanged with 2 errors, and increased with 3 or more errors, maintaining average performance between 66% and 83% correct across orientations. Participants with output threshold larger than 10% were excluded from further analysis. **Practice:** Participants first completed a two-stage practice. Stage 1 consisted of 8 trials with progressive difficulty, starting at 40% with a step factor of 0.3 log units. Stage 2 consisted of 20 trials following a 3-down-1-up staircase. For all participants, the starting contrast was set to 10%. **Main task:** The starting contrast for each participant was set to the smaller of 10% contrast or twice the threshold obtained during practice, to mitigate cold-start performance effects. The outcome measure was the contrast threshold (% Michelson contrast), defined as the average contrast level across the last 10 trials.

**Figure 1.**
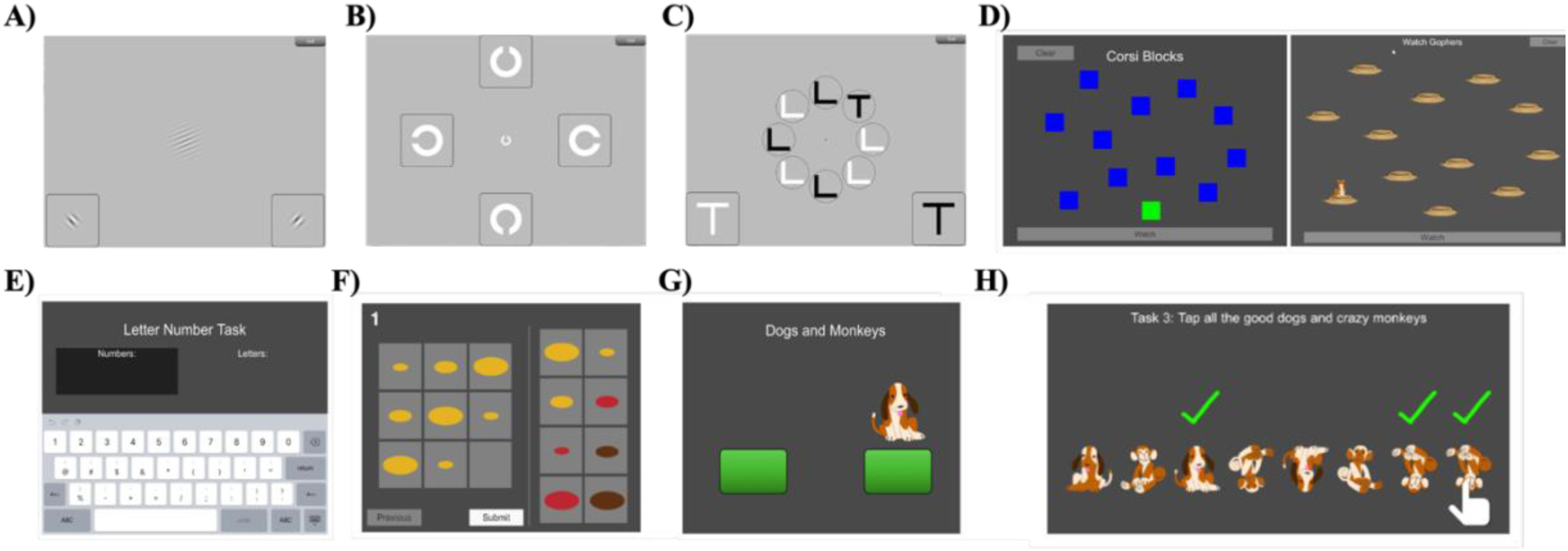
Illustration of the ten tasks administered in the study. Visual assessment tasks: A) Gabor patch stimulus used for the Contrast Sensitivity Function, Spatial Frequency Cutoff, and CSF Cutoff tasks B) Sloan letter C used in the Visual Acuity task C) Visual Search display. Cognitive assessment tasks: (D) Simple Corsi E) Letter Number sequencing display F) University of California Matrix Reasoning Task (UCMRT) G) Countermanding task H) UCancellation task.

#### 2.5.2. Visual Acuity (VA)

Visual acuity was assessed using a block letter C (Sloan Font) presented in one of four orientations (gap facing up, down, left, or right). This stimulus was chosen because it provides a standardized measure of spatial resolution and is widely used as a reference optotype in clinical and psychophysical assessments of visual acuity (Bailey & Lovie, 1976; Ferris et al., 1982). Participants indicated the direction of the gap by pressing the corresponding C icon placed in one of the four cardinal directions on the display (Figure 1B). Participants with outputs larger than the starting value were excluded. **Practice:** Participants completed 12 trials with progressive difficulty (1.3, 1.1, 0.9, 0.7, 0.5, 0.4, 0.3, 0.2, 0.1, 0.0, −0.1, and −0.2 logMAR units). **Main task:** Following practice, participants completed the main task using a conventional 3-down-1-up staircase terminating after 60 trials. The starting stimulus size was 0.5 logMAR (0.2634 deg of visual angle). The staircase proceeded in two stages: a step factor of 0.2 log units was used until 3 reversals (Stage 1), after which the step factor was reduced to 0.1 log units (Stage 2) until task completion. The outcome measure was the acuity threshold in degree of visual angle.

#### 2.5.3. Spatial Frequency (SF) Cutoff

The SF cutoff task measured the highest spatial frequency resolvable at a fixed contrast level, using a 2-degree Gabor patch. Participants indicated the direction of tilt (45° or 135°) by pressing the corresponding icon on the display (Figure 1A). Stimulus contrast was fixed at 12.5%. For this task, we use a cutoff exclusion criterion of 10 cpd. **Practice:** A two-stage practice was administered. Stage 1 consisted of 8 trials following a 1-up-1-down staircase with a starting spatial frequency of 6 cpd. Stage 2 consisted of 20 trials with a starting frequency of 10 cpd, also following a simple staircase. **Main task:** The main task employed a 3-down-1-up staircase terminating after 60 trials, with a starting spatial frequency of 9 cpd. The outcome measure was the SF cutoff threshold in cpd.

#### 2.5.4. CSF Cutoff

The CSF cutoff task estimated contrast sensitivity at or near each participant’s individual spatial frequency cutoff, i.e., the most demanding spatial conditions for their visual system. A 1-degree Gabor patch was used, and the spatial frequency was set individually based on each participant’s SF cutoff threshold obtained in Section 2.5.3. Participants indicated the direction of tilt either 45° or 135° (Figure 1A). Participants with output thresholds larger than 30% were excluded from the study. **Practice:** Stage 1 consisted of 8 trials with progressive reduction of stimulus contrast. Stage 2 consisted of 20 trials following a 3-down-1-up staircase, with a starting contrast of 50% (−0.3 log units). **Main task:** The main task followed the same blockwise staircase structure as the CSF task (20 mini-blocks of 6 trials, 6 orientations per block). The initial stimulus contrast was set to the minimum of 50% or twice the participant’s practice threshold. The outcome measure was the contrast threshold at the individual SF cutoff (logCS).

#### 2.5.5. Visual Search

The visual search task assessed the ability to rapidly identify a target among distractors under increasing temporal demands. Participants reported the color (black or white) of a target element—a right-side-up “T”––by pressing the corresponding response button (Figure 1C). The target was presented among distractor elements consisting of upside-down “L” shapes in both black and white. Participants were instructed to ignore distractors and identify the target color as quickly and accurately as possible. **Practice:** Stage 1 comprised 8 trials with progressively reducing stimulus-to-mask intervals (from 600 ms to 30 ms in 50 ms steps). Stage 2 comprised 20 trials following a 2-up-1-down staircase, starting at 500 ms with 8.33 ms steps. **Main task:** The main task’s stimulus-to-mask interval started at the final value of the practice staircase and employed a 3-down-1-up staircase terminating after 60 trials. The staircase proceeded in two stages: a step of 25 ms until 3 reversals (Stage 1), after which the step was reduced to 8 ms (Stage 2). The outcome measure was the stimulus duration threshold in milliseconds.

### 2.6. Cognitive assessment tasks

All cognitive tasks were administered via the Recollect Assessment Battery (RAB), a tablet-based battery developed at the UCR Brain Game Center that runs on iOS and Android (Pahor, Seitz, et al., 2022). Tasks were delivered through the PLFest application and the Recollect app across sites. Similar to visual assessment tasks, before conducting the experiment, all cognitive tasks include standardized instructions and practice sessions.

#### 2.6.1. Visuospatial working memory––Simple Corsi Task

Visuospatial working memory was assessed using the tablet-based adaptation of the traditional Corsi Block-Tapping Test (Brunetti et al., 2014). Participants viewed a 4×4 grid of 12 spatially distributed squares, a subset of which illuminated sequentially to form a spatial sequence (Figure 1D). Following the presentation of the full sequence, participants reproduced it by tapping the squares in the same order. Set size—defined as the number of locations in the sequence—increased progressively across trials, with the starting set size of 2 increasing as performance criteria were met. The Simple Corsi task follows the classic “method of limits” approach, in which 2 trials are presented per set size and the task terminates when both trials at a given set size are answered incorrectly. **Practice:** Participants completed 2 practice trials at set sizes 2 and 3 before the main task. **Main task:** The primary outcome measure was *span*, defined as the highest set size at which at least one trial was answered correctly. Sequences were drawn randomly without repetition from a pre-generated pool of 12 sequences per set size (set sizes 2–10), making the task suitable for repeated testing.

#### 2.6.2. Verbal working memory––Letter-Number Task

Verbal working memory was assessed using the Letter-Number Task, a subtest of the Wechsler Adult Intelligence Scale (WAIS) (Valentine et al., 2020). Participants were presented with a mixed sequence of letters and numbers and were required to mentally reorder them, reporting the numbers in ascending order followed by the letters in alphabetical order (Figure 1E). For example, the sequence “H8T3K5” would be reordered to “358” and “HKT.” Set size (sequence length) ranged from 2 to 15. **Practice:** Participants completed a 5-trial practice session (2 trials at set size 2, 3 trials at set size 3). At least 3 out of 5 practice trials had to be correct to proceed, otherwise, practice was repeated once. **Main task:** The task followed a method of limits procedure in which 2 trials were presented per set size in ascending order of difficulty. The task terminated when both trials at a given set size were answered incorrectly (stopping rule). Primary outcome measure was *span*––the highest set size at which at least one trial was answered correctly.

#### 2.6.3. Fluid intelligence––Matrix Reasoning (UCMRT)

Fluid intelligence and abstract problem-solving were assessed using the University of California Matrix Reasoning Task (UCMRT), a validated tablet-based measure with three alternate forms designed for college-level populations (Pahor et al., 2019). UCMRT consists of Raven-like matrix problems in which participants identify the missing element in a 3×3 matrix from 8 answer alternatives (Figure 1F). Problems were developed from matrices produced by Sandia National Laboratories (Matzen et al., 2010) and redesigned with larger, non-overlapping stimuli in color to ensure clarity on tablet displays. **Practice:** Participants received feedback (correct/incorrect) and explicit rule explanations during practice. A minimum accuracy criterion of 4 out of 6 correct was required to advance to the test phase; if not met, practice was repeated with an alternate problem set. **Main task:** The 23 test problems were completed within a 10-minute time limit. No feedback was provided during the test, but participants could change their answers, skip problems, and navigate freely between items. The primary outcome measure was the *total number of correctly solved problems*. UCMRT has demonstrated strong convergent validity with Raven’s Advanced Progressive Matrices (*r* =.58, *N* = 233; Pahor et al., 2019) and predictive validity for academic performance in college samples.

#### 2.6.4. Inhibitory control––Countermanding

Inhibitory control and executive function were assessed using the Countermanding task. The task requires participants to tap one of two green buttons in response to a visual stimulus—a picture of a dog or a monkey—presented on the left or right side of the screen (Figure 1G). For congruent trials (dog), participants must tap the button on the same side as the stimulus. For incongruent trials (monkey), participants must suppress the prepotent response to respond on the same side and instead tap the button on the opposite side. **Practice:** A separate practice block preceded each of the three test blocks (congruent, incongruent, and mixed), providing participants with experience of each trial type before the test phase. **Main task:** The task consisted of three blocks: a congruent block (12 trials), an incongruent block (12 trials), and a randomized mixed block (48 trials) in which congruent and incongruent trials were intermixed. In the mixed block, 50% of trials were switch trials, with no more than 2 consecutive switch or non-switch trials. The primary outcome measure were mean reaction times for correct responses in the mixed block for congruent trials (index of Processing Speed) and incongruent trials (index of Inhibitory Control).

#### 2.6.5. Selective attention––Ucancellation task

UCancellation is a gamified, tablet-based cancellation task modeled after the D2 Test of Attention, validated for mobile delivery by (Pahor, Mester, et al., 2022). Rather than letters, the task uses cartoon images of dogs and monkeys as stimuli, enhancing engagement and suitability for diverse populations, including children. Each row displays 8 items containing 3–5 targets. Participants must select targets—upright dogs (tail on the left) and upside-down monkeys (tail on right)—separately in single-target blocks and together in a mixed block where both target types are present simultaneously. The global time limit is 3 minutes and 30 seconds for the mixed block. The task is designed to measure selective attention, inhibitory control, and concentration. **Practice:** Participants completed a tutorial followed by a practice session for each target type (dogs, then monkeys) consisting of 2–3 rows with performance feedback. A mixed-block practice of 3 rows preceded the test block. **Main task:** The mixed block constituted the primary assessment phase, with a minimum of 30 rows and 120 targets. The primary outcome measure was *Concentration Performance*, calculated as ΣHits − ΣFalse Alarms in the mixed block, consistent with Pahor et al. (2022).

### 2.7. Statistical Analysis

Descriptive statistics are shown for all visual and cognitive measures across the full sample. Normality was assessed using the Shapiro-Wilk test. To test for differences in visual assessment performance across testing sites, we conducted a Bayesian one-way ANOVA for each of the five visual tasks, using site as the between-subjects factor and mean threshold as the dependent variable, implemented via the *BayesFactor* package in R with a Cauchy prior scale of *r* = 0.5. We report BF₀₁, such that larger values indicate stronger evidence in favor of the null hypothesis of cross-site equivalence. We interpreted BF₀₁ according to established conventions, where BF₀₁ > 3 indicates moderate to strong evidence for cross-site equivalence.

## 3. Results

### 3.1. Visual assessments

Figure 2 presents, for each of the five visual assessment tasks, the trial-by-trial threshold trajectories across the full task sequence. Individual participants are shown as thin colored lines, and the group mean is overlaid as a thick line. Additionally, Figure 3 shows violin plots for each of the five visual assessments displaying the distribution of the mean threshold estimates across sites and participants. The right-side panels show the breakdown of those distributions by testing site (NU, UCR, UW, and Mills) with bar height indicating the site mean and error bars reflecting the standard error of the mean (SEM). Together, these figures serve complementary purposes: the trajectory plots demonstrate that PLFest’s adaptive staircase procedures converge reliably to stable threshold estimates, the violin plots confirm that the resulting group distributions are consistent with normative values reported in the literature and the bar plot illustrate the cross-site robustness of PLFest’s visual assessments across the four testing locations.

#### 3.1.1. Visual Acuity

Trial-by-trial threshold trajectories for the VA task (expressed in logMAR) are shown in Figure 2A. Individual thin lines revealed the characteristic two-phase convergence pattern produced by the two-stage staircase: a steeper, coarser descent during Stage 1 (step size 0.2 logMAR, until 3 reversals), followed by finer oscillation around the estimated threshold during Stage 2 (step size 0.1 logMAR, remaining 60 trials). The group mean trajectory stabilized at approximately-0.06 logMAR by trial 20, consistent with near-normal acuity in our sample and recent studies estimating population-level visual acuity (Peñaloza et al., 2025; Peñaloza & Kwon, 2025) using clinical eye-charts (dashed horizontal line in Figure 3A).

**Figure 2.**
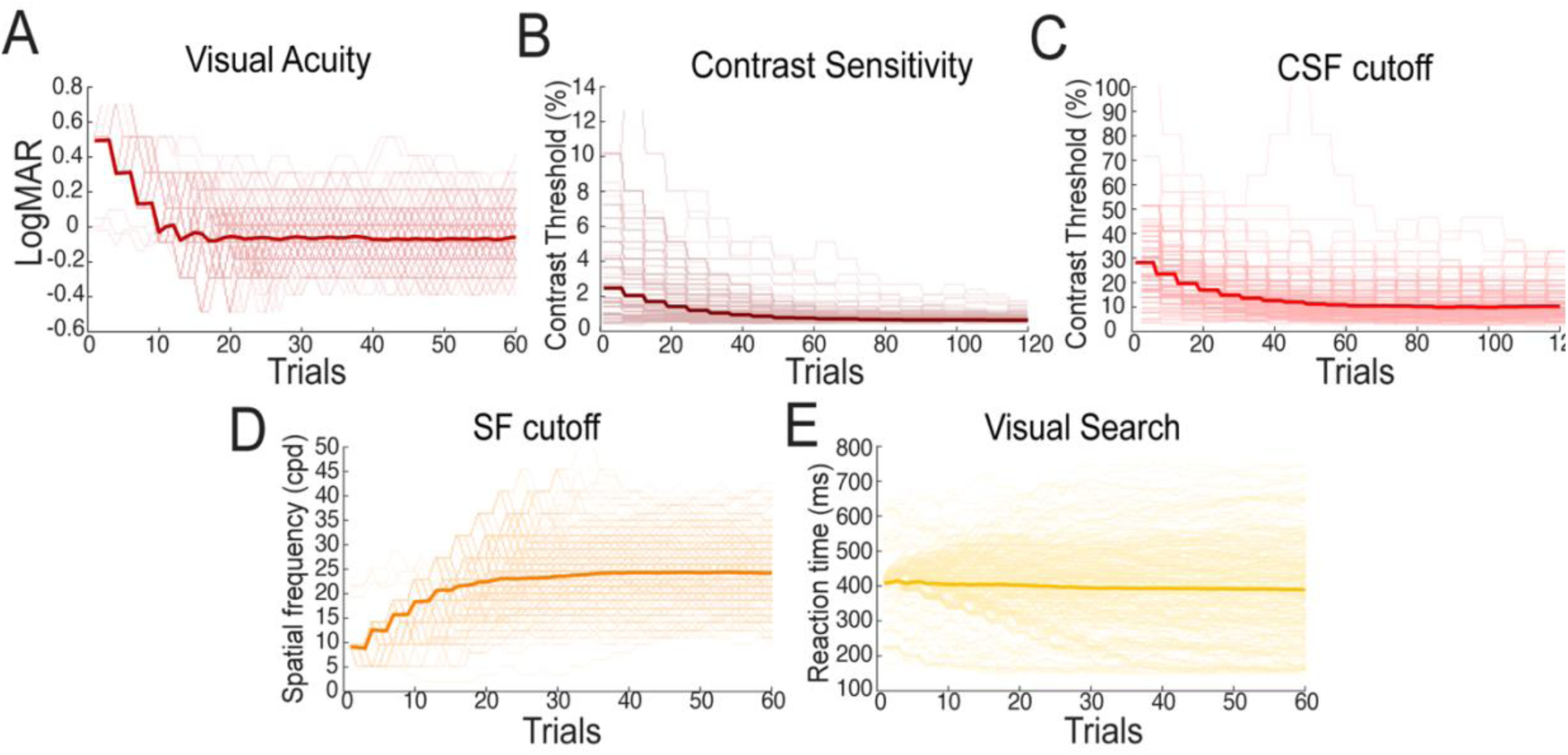
Trial-by-trial staircase trajectories for the five visual assessment tasks. Individual participant staircases are shown as thin colored lines and the group mean as a thick line. Panels show: (A) visual acuity (logMAR), (B) contrast sensitivity threshold (% Michelson contrast), (C) CSF cutoff contrast threshold (%), (D) spatial frequency cutoff (cpd), (E) visual search reaction time (ms), and (E) Visual search. Across most tasks, the group mean trajectory converges rapidly toward a stable threshold estimate, demonstrating the reliability of PLFest’s adaptive staircase procedures.

**Figure 3.**
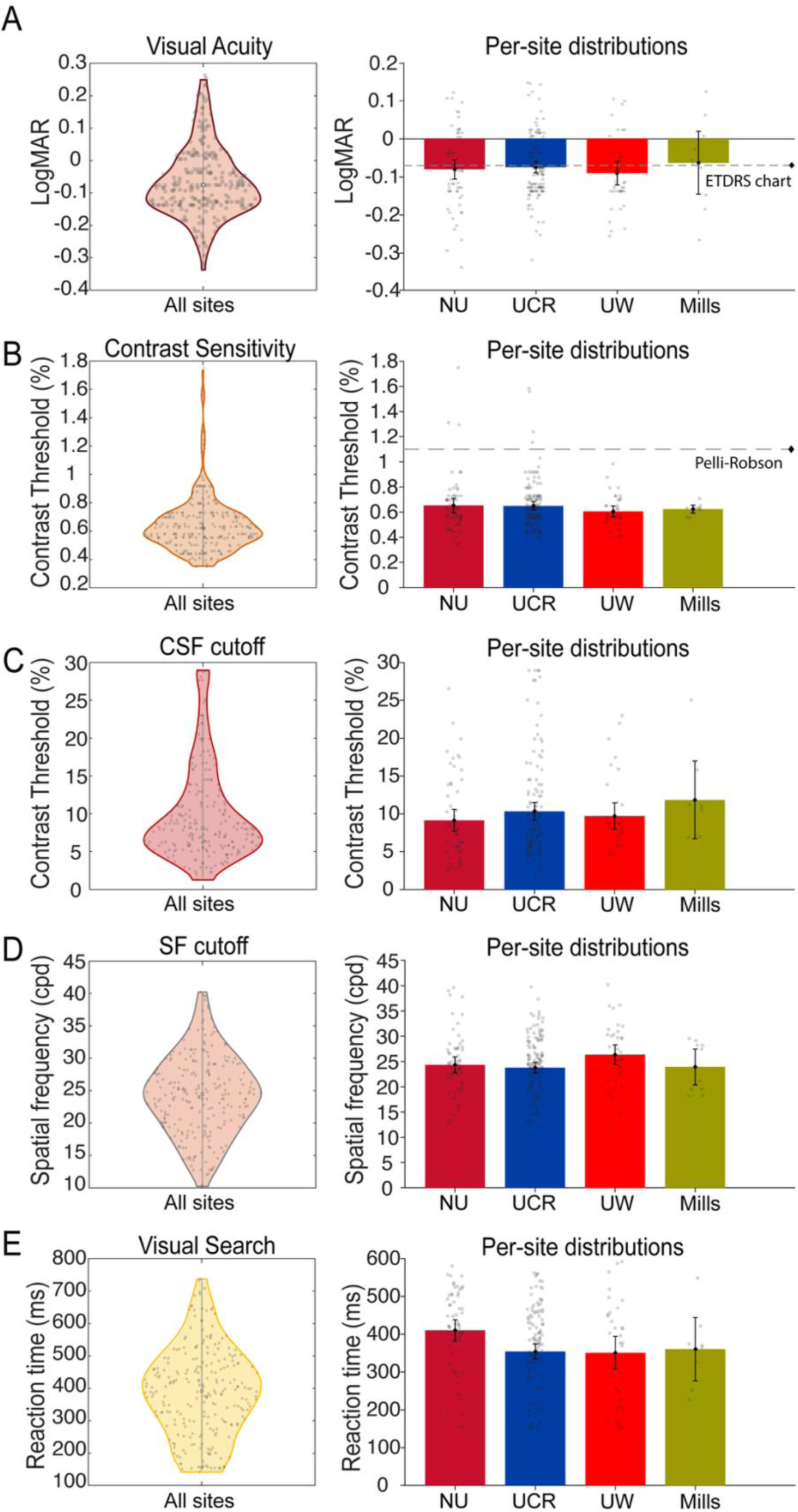
Performance distributions for the five visual assessment tasks across the full sample and by testing site. For each task, the left panel displays a violin plot of the distribution of mean threshold estimates across all participants, with individual data points jittered within; the right panel shows per-site mean performance (±SEM) for the four testing locations (NU, UCR, UW, Mills), with individual participant scores overlaid as grey dots. Reference lines in panels A and B indicate normative benchmarks from the ETDRS chart (−0.07 logMAR) and Pelli-Robson chart (∼1.0% contrast threshold), respectively. Panels show: (A) visual acuity (logMAR), (B) contrast sensitivity threshold (%), (C) CSF cutoff contrast threshold (%), (D) spatial frequency cutoff (cpd), and (E) visual search reaction time (ms).

**Figure 3.**
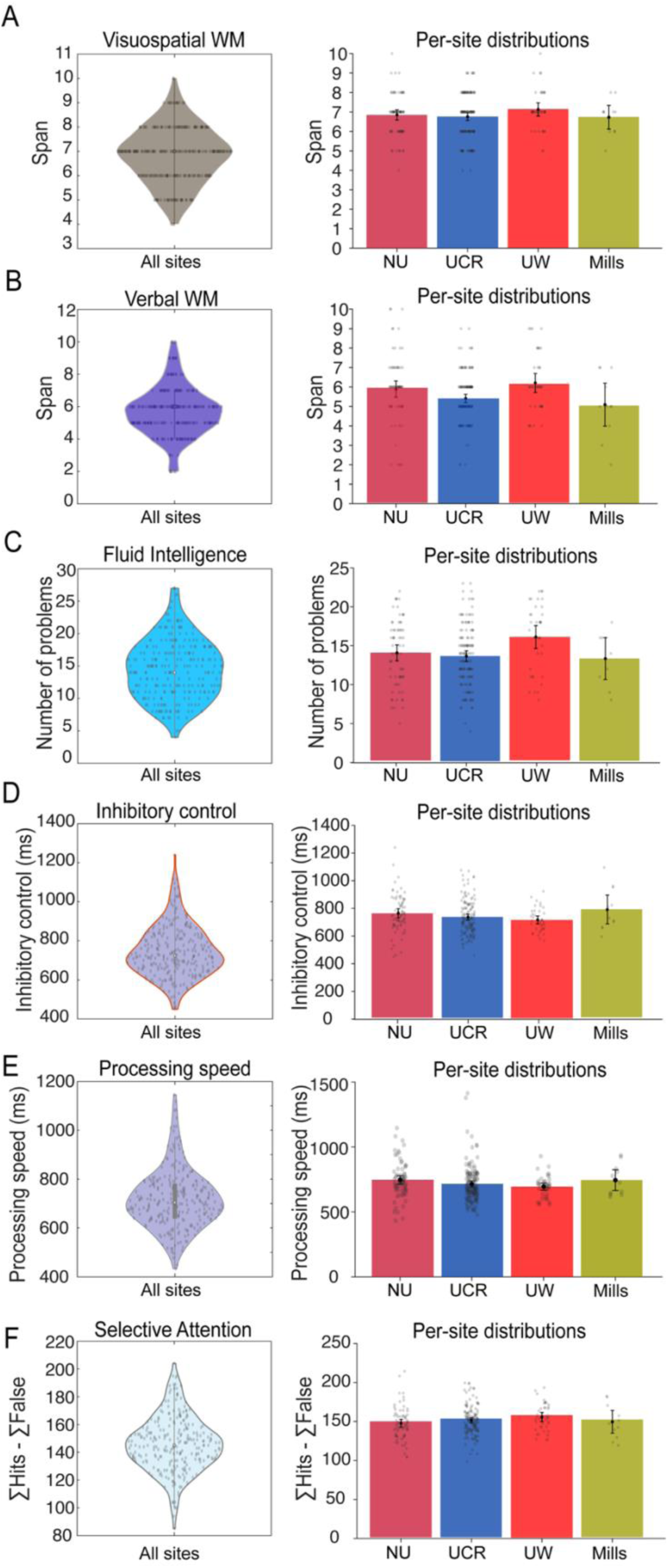
Performance distributions for the five cognitive assessment measures across the full sample and by testing site. For each measure, the left panel displays a violin plot of the distribution of scores across all participants. The right panel shows per-site mean performance (±SEM) for the four testing locations. Panels show: (A) visuospatial working memory span (Simple Corsi), (B) verbal working memory span (Letter-Number Task), (C) fluid intelligence — number of correctly solved problems (Matrix Reasoning), (D) inhibitory control reaction time in ms (Countermanding), (E) processing speed reaction time in ms (Countermanding), and (F) selective attention concentration performance score (ΣHits − ΣFalse Alarms; UCancellation).

The violin plot in Figure 3A shows the distribution of mean acuity thresholds (M =-0.07, SD = 0.1 logMAR; Snellen equivalent ≈ 20/16). The distribution was moderately right-skewed (skew=0.65; Shapiro-Wilk, p < 0.05) and consistent with the expected range for a young adult sample with normal or corrected-to-normal vision (Peñaloza et al., 2025; Peñaloza & Kwon, 2025). These values are in good agreement with tablet-based VA norms reported by Jayakumar et al. (2024) and the web-based DDART validation study (Labiris et al., 2023). A Bayesian one-way ANOVA revealed moderate evidence for cross-site equivalence in visual acuity thresholds (BF₀₁ = 7.25), supporting the robustness of PLFest’s VA assessment across testing locations.

#### 3.1.2. Contrast Sensitivity Function

As shown in Figure 2B, individual threshold trajectories for the CSF task (expressed in % Michelson contrast) showed rapid initial adjustment from each participant’s starting contrast level, followed by increasingly tight oscillation around a stable estimate as the task progressed across the 120 trials. While individual thin lines reflected the expected inter-subject variability in absolute sensitivity, the trajectories remained well-bounded and did not show systematic drift across trials, indicating that the blockwise staircase procedure successfully and continuously tracked each participant’s contrast threshold. The group mean trajectory (thick line) descended smoothly from the starting value and stabilized at approximately 0.7% by trial 50, with only small fluctuations after that.

Figure 3B shows the distribution of mean contrast thresholds across participants (M = 0.6%, SD = 0.2%). The distribution showed that most participants performed within a well-defined sensitivity range, with a few observers having larger thresholds right-skewing the distribution (skew=2.25; Shapiro-Wilk, p < 0.05). These values are consistent with contrast sensitivity norms for healthy young adults measured on tablet-based devices (Habtamu et al., 2019; Jayakumar et al., 2024; Kollbaum et al., 2014) and yielded finer resolution in threshold estimation compared to the discrete scoring increments of standard clinical charts such as the Pelli-Robson (dashed line in Figure 3B). Finally, a Bayesian one-way ANOVA provided strong evidence that PLFest yields consistent contrast sensitivity estimates regardless of testing location (BF₀₁ = 12.0).

#### 3.1.3. Spatial frequency cutoff

Individual threshold trajectories for the SF cutoff task (expressed in cpd) are shown in Figure 2D. Trajectories increased from the 9 cpd starting value across the 60 trials, with individual thin lines showing somewhat greater variability than observed for the CSF or VA tasks. This is expected, as the SF cutoff probes the upper limit of spatial resolution at a fixed, relatively low contrast (12.5%), a region of the contrast sensitivity function where inter-individual differences are more pronounced. The group mean trajectory converged to approximately 23 cpd by trial 20.

The violin plot displays the distribution of SF cutoff thresholds (M = 24.32 cpd, SD = 6.04 cpd). The distribution was normally distributed (skew=0.13; Shapiro-Wilk, p = 0.25), with most participants falling within the range of 10 and 40 cpd, consistent with normal spatial resolution limits in healthy young adults (Campbell et al., 1978; Campbell & Robson, 1968). Importantly, the observed mean SF cutoff of 24.32 cpd is lower than the expected grating acuity limit of ∼30 cpd for 20/20 vision, likely because the fixed 12.5% contrast intersects the descending high-frequency limb of the CSF before the absolute resolution limit is reached (Campbell & Robson, 1968; Watson & Ahumada, 2005). Finally, the Bayesian analysis indicated that SF cutoff estimates are highly consistent across testing locations (BF₀₁ = 10.4).

#### 3.1.4. CSF cutoff

The CSF cutoff task estimated contrast sensitivity at each participant’s individual SF cutoff frequency. As a consequence, the starting trial value and absolute threshold level differed across participants, as shown in the spread of individual thin-line trajectories in Figure 2C. Nevertheless, within each participant the trajectory showed clear convergence, and the group mean trajectory stabilized by approximately trial 60.

The mean CSF cutoff threshold across participants was M = 10.1%, SD = 6.34%. similar to the CSF data, the CSF cutoff distribution showed right-skewness (skew=1.34; Shapiro-Wilk, p < 0.05), reflecting the variability in the individual spatial frequency cutoffs. The Bayesian one-way ANOVA showed moderate support for cross-site equivalence in CSF cutoff thresholds (BF₀₁ = 3.0).

#### 3.1.5. Visual Search

In the visual search task, threshold trajectories reflected the minimum stimulus presentation duration (in ms) at which participants could reliably identify the target among distractors. As shown in Figure 2E, there was significantly greater inter-individual variability in threshold duration compared to the other visual assessments, as evidenced by the wider spread of individual staircase trajectories across trials.

Visual search threshold mean and standard deviation were M = 368.97 and SD = 115.91 ms, respectively. The distribution was slightly left-skewed (skew=-0.24; Shapiro-Wilk, p < 0.05) with most participants achieving thresholds in the range between 142 and 592 ms. The large inter-individual variability in visual search thresholds is consistent with the well-established dependence of visual search on attention guidance, target-distractor similarity, and individual differences in attentional deployment (Wolfe & Horowitz, 2017), all of which may affect performance on individual subjects. In line with this inter-individual variability, a Bayesian one-way ANOVA did not yield evidence for cross-site equivalence in visual search reaction times (BF₀₁ = 0.13).

### 3.2. Cognitive assessments

Figure 4 presents violin plots of the distributions of performance scores for the five cognitive assessment tasks administered in our study as well as bar plots with the breakdown of participants across testing sites.

#### 3.2.1. Simple Corsi

The distribution of visuospatial working-memory span scores yielded a mean span of M = 7, SD

= 1. The distribution was slightly left-skewed (skew=-0.04; Shapiro-Wilk, p < 0.05), with most participants achieving spans in the 4 to 10 range. These values are consistent with other sources reporting data for tablet-based adaptations of the Corsi Block-Tapping Test in young adult (Brunetti et al., 2014). The absence of pronounced floor or ceiling effects confirms that the Simple Corsi task, as delivered through our application, is sensitive to individual differences in visuospatial short-term storage capacity across this population. Reinforcing this cross-site consistency, a Bayesian one-way ANOVA provided strong evidence for equivalence in visuospatial working memory span across testing sites (BF₀₁ = 10.5).

#### 3.2.2. Letter Number Task

Verbal working memory span yielded a mean of M = 6, SD = 2. The distribution was slightly right-skewed (skew= 0.23; Shapiro-Wilk, p < 0.05) with scores ranging from 2 to 10. These values are consistent with values reported for the Letter-Number Sequencing subtest of the Wechsler Adult Intelligence Scale (WAIS) in college-aged samples. Like in the Simple Corsi task the spread of span scores across participants indicates that the task is sensitive enough to capture individual variability across the range of verbal working memory capacity. Additionally, the observed span values were comparable to those obtained on the Simple Corsi task, consistent with the general pattern of comparable capacity limits across verbal and visuospatial working memory domains documented in the literature (Baddeley, 2003; Cowan, 2001). In contrast to the visuospatial span measure, however, a Bayesian one-way ANOVA did not yield evidence for cross-site equivalence in verbal working memory span (BF₀₁ = 0.22). This result may stem form the high inter-individual variability inherent to span measures, reducing the sensitivity to detect equivalence.

#### 3.2.3. Matrix Reasoning

Performance on the UCMRT yielded a mean accuracy of M = 14 correct items (SD = 6) out of 23 problems within the 10-minute time limit. The distribution was moderately right-skewed (skew= 0.31; Shapiro-Wilk, p < 0.05). These values are in close agreement with normative data from a large-scale validation study of the UCMRT in college-aged samples (Pahor et al., 2019), which reported strong convergent validity with Raven’s Advanced Progressive Matrices (*r* =.58, *N* = 233) and predictive validity for academic performance. The convergence of our sample’s performance with the established normative benchmark for this task provides direct evidence that the present multi-site sample is representative of the population for which the task was designed. Despite this normative alignment, a Bayesian one-way ANOVA yielded only anecdotal evidence for cross-site equivalence in matrix reasoning performance (BF₀₁ = 0.85), indicating that the data are presently inconclusive with respect to site consistency and that replication with larger samples is needed.

#### 3.2.4. Countermanding

The Countermanding task yielded mean reaction times for correct responses in the mixed block of M = 716.78 ms (SD = 120.57 ms) for Processing Speed (congruent trials) and M = 745.33 ms (SD = 128.88 ms) for Inhibitory Control (incongruent trials). Distributions were right-skewed for both conditions (Shapiro-Wilk, p < 0.5; skew = 0.73, 0.63 for Processing Speed and Inhibitory Control, respectively), consistent with the positive skew typically observed in reaction time distributions. Accuracy was very high across all trial types (overall accuracy = 95%), as expected in a healthy young adult sample, confirming that reaction time rather than accuracy is the informative dependent measure in this task. These results align closely with the normative data reported by Pahor et al. (2022), who observed mean reaction times of 804.28 ms (SD = 112.74) for Processing Speed, and 822.02 ms (SD = 113.43) for Inhibitory Control in a college-aged sample using the same task implementation. The close correspondence between our multi-site sample and these benchmarks shows that our study delivers high fidelity data across distributed testing sites. In terms of multi-site consistency, a Bayesian one-way ANOVA provided strong evidence for cross-site equivalence in Countermanding performance (BF₀₁ = 16.0), indicating that inhibitory control and processing speed estimates derived from this task are highly robust across distributed testing locations.

#### 3.2.5. UCancellation

Selective attention and inhibitory control, as measured by the UCancellation task, yielded a mean concentration performance of M = 147.02 (SD = 19.49). The distribution was normally distributed with slight right-skewness (skew= 0.20; Shapiro-Wilk, p = 0.13), consistent with the moderate positive skew in CP documented in Pahor et al., (2022). Our values are comparable to the normative benchmark reported by Pahor et al. (2022) for UCancellation Pictures (M = 143.04, SD = 20.73). The convergence of UCancellation performance in the present study with previously established norms provides evidence that the gamified picture-based implementation of the cancellation paradigm yields valid assessments of selective attention and cognitive control in a large multi-site context. Corroborating this interpretation, a Bayesian one-way ANOVA provided strong evidence for cross-site equivalence in UCancellation concentration performance (BF₀₁ = 15.0), further supporting the reliability of our platform to collect data across geographically distributed testing sites.

**Table 1.**
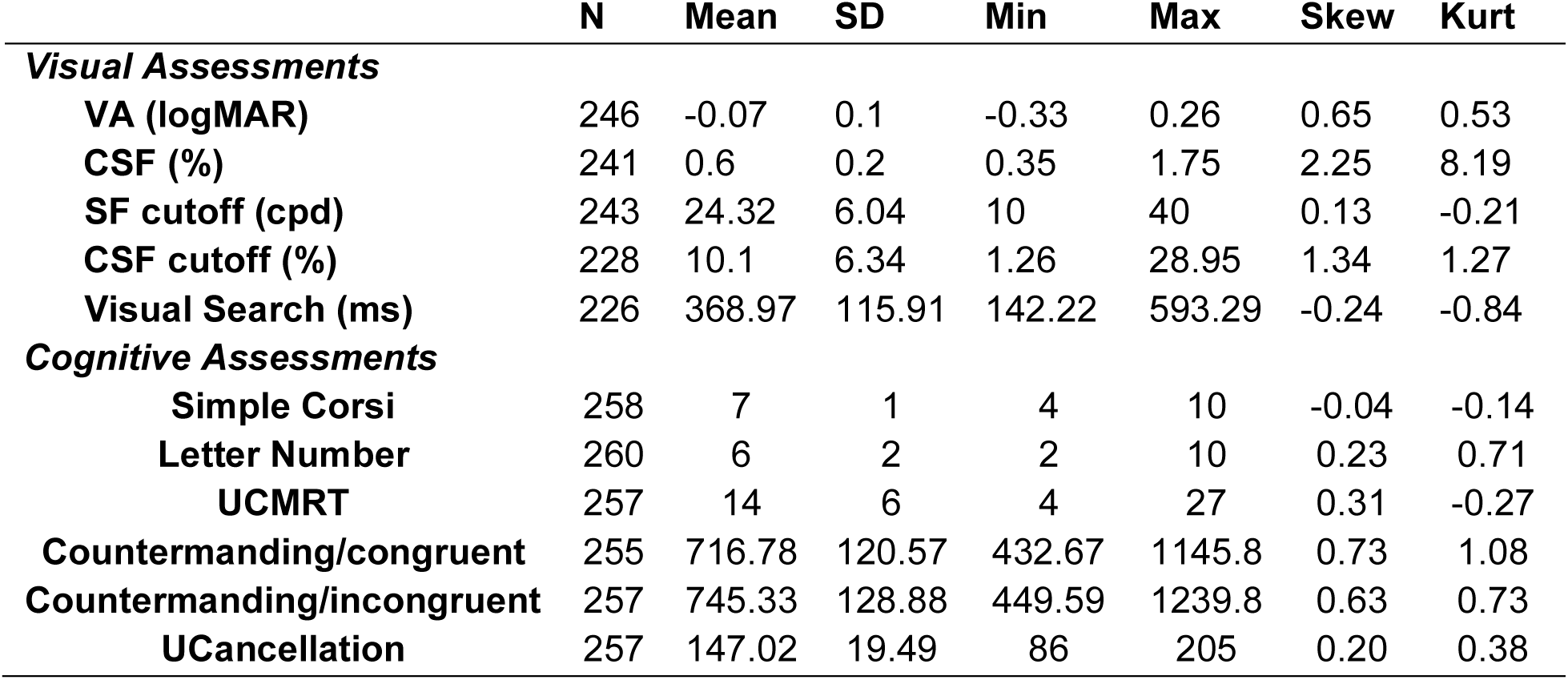
Summary of descriptive statistics for all ten assessments.

## 4. Discussion

Vision science is currently undergoing a major transformation in which assessment and training paradigms, once confined to laboratory environments requiring highly specialized equipment, are increasingly being translated into portable and scalable digital tools with applications in both research and clinical practice. Over the past decade, this shift has driven the development of digital psychophysical platforms capable of collecting perceptual and cognitive data outside traditional laboratory settings. Recent studies have demonstrated that tablet-and smartphone-based measures of fundamental visual functions, including contrast sensitivity (CS) and visual acuity (VA), can achieve performance comparable to laboratory and clinical standards while maintaining acceptable reliability (Habtamu et al., 2019; Kollbaum et al., 2014; Labiris et al., 2023). However, most existing platforms remain focused on isolated assessment domains, limiting their utility for comprehensive evaluation of perceptual and cognitive function.

To address these limitations, we developed PLFest, a cross-platform application designed to support the measurement and training of a broad range of perceptual and cognitive functions within a unified framework (Jayakumar et al., 2024). Building on our initial validation of PLFest for assessments of visual acuity and contrast sensitivity, the present study extends this work by reporting baseline data from a large ongoing multi-site clinical trial involving participants recruited across four geographically distributed university sites. The primary objective was to determine whether PLFest could provide reliable and valid measurements across a diverse battery of visual and cognitive tasks under distributed testing conditions.

Overall, the findings support the validity and scalability of the platform. Across all tasks, the group mean trajectory converges rapidly toward a stable threshold estimate, demonstrating the reliability of PLFest’s adaptive staircase procedures. The distributions of visual acuity and contrast sensitivity closely matched population-level normative data reported in previous psychophysical and clinical studies (Peñaloza et al., 2025; Peñaloza & Kwon, 2025). In particular, average visual acuity values were highly consistent with normative performance reported in healthy young adults tested using standardized chart-based methods (Elliot et al., 1995; Peñaloza et al., 2025; Peñaloza & Kwon, 2025). Similarly, contrast sensitivity thresholds closely aligned with tablet-based and laboratory-based normative datasets reported by Kollbaum et al. (2014) and Habtamu et al. (2019). Performance across the cognitive assessment battery also demonstrated good agreement with previous studies from our group employing the same paradigms (Pahor et al., 2019, 2022). Together, these convergent findings provide strong evidence that PLFest can generate high-quality visual and cognitive measurements using standardized consumer-grade hardware.

Although most tasks produced stable and well-distributed performance measures, some paradigms showed greater variability than others. Visual search performance exhibited substantial inter-individual variability, and some participants performing the spatial frequency cutoff task reached thresholds above the initial starting values of the adaptive procedure. These findings suggest that, in addition to genuine individual differences, some participants may have experienced difficulties understanding or performing specific tasks. While all participants completed practice phases prior to testing, future implementations may benefit from more detailed instructions or additional practice trials to further improve measurement reliability. In the case of the spatial frequency cutoff task, the target contrast level may also have been overly challenging for some individuals, potentially contributing to the lower average performance observed relative to classical normative estimates of about 30 to 60 cpd (Campbell & Robson, 1968; Watson & Ahumada, 2005).

Importantly, the elevated variability observed in the visual search task may represent an informative feature rather than simply a methodological limitation. Unlike visual acuity and contrast sensitivity, which primarily reflect relatively low-level sensory processing, visual search performance depends heavily on higher-order cognitive processes, including attentional allocation, perceptual organization, target-distractor similarity, and processing speed (Wolfe & Horowitz, 2017). The broader cognitive demands of this task likely contributed to the wider distribution of performance observed in our sample and suggest that visual search may serve as a particularly sensitive indicator of individual differences in attentional and cognitive functioning within the PLFest battery.

Despite these opportunities for refinement, the overall pattern of results demonstrates that PLFest provides robust measurements across a diverse range of perceptual and cognitive domains. Performance distributions were generally well-behaved and closely aligned with established normative benchmarks, indicating that high-quality psychophysical and cognitive data can be collected using standardized consumer-grade hardware outside traditional laboratory settings. Importantly, these findings were observed across four geographically distributed sites, supporting the platform’s suitability for multi-site research and large-scale data collection efforts. Together, the results provide evidence that portable digital platforms can maintain the methodological rigor required for psychophysical assessment while substantially increasing accessibility and scalability.

The ability to integrate perceptual and cognitive assessments within a unified digital environment creates important opportunities for future research. Although perceptual learning and cognitive training paradigms have often developed independently, growing evidence suggests that perceptual learning interacts with broader cognitive systems including attention, working memory, and executive control (Kannan et al., 2024). Platforms such as PLFest provide a scalable and standardized framework for investigating these interactions across diverse populations and testing environments. Furthermore, by reducing dependence on specialized laboratory infrastructure, PLFest may facilitate broader participation in research and support the development of larger, more representative normative datasets.

Several limitations should be acknowledged. First, the present analyses were conducted on baseline data from a predominantly university-based sample and therefore may not fully capture the variability present in the general population. Second, although testing procedures were standardized across sites, some environmental variability is unavoidable in distributed testing contexts. Third, the current study focused on healthy participants; additional validation in clinical populations will be necessary to establish the platform’s utility for diagnosis, monitoring, and rehabilitation. Future work should also examine longitudinal reliability and sensitivity to training-related changes, particularly in populations with visual or cognitive impairments.

In summary, recent advances in digital technology have created new opportunities to assess visual and cognitive function using accessible, portable, and user-friendly platforms that integrate multiple assessments within a single device. Across a large multi-site sample, PLFest produced measures of visual acuity, contrast sensitivity, and cognitive performance that were consistent with established normative benchmarks, supporting its validity as a scalable assessment tool. By automating administration, scoring, and data management, the platform reduces reliance on specialized equipment and highly trained personnel while maintaining the precision required for psychophysical research. These capabilities position PLFest as a valuable resource for large-scale studies of perceptual learning, cognitive function, and brain plasticity, as well as for future clinical applications in screening, monitoring, and rehabilitation. More broadly, standardized digital platforms such as PLFest may help address longstanding challenges of reproducibility, accessibility, and scalability that have historically limited both basic and translational vision science.

## References

Aberg, K. C., & Herzog, M. H. (2010). Does perceptual learning suffer from retrograde interference? PLoS ONE, 5(12), e14161. 10.1371/journal.pone.0014161

Baddeley, A. (2003). Working memory: looking back and looking forward. Nature Reviews Neuroscience, 4(10), 829–839. 10.1038/nrn1201

Bailey, I. L., & Lovie, J. E. (1976). New design principles for visual acuity letter charts. Am J Optom Physiol Opt, 53(11), 740–745. 10.1097/00006324-197611000-00006

Bastawrous, A., Rono, H. K., Livingstone, I. A., Weiss, H. A., Jordan, S., Kuper, H., & Burton, M. J. (2015). Development and Validation of a Smartphone-Based Visual Acuity Test (Peek Acuity) for Clinical Practice and Community-Based Fieldwork. JAMA Ophthalmol, 133(8), 930–937. 10.1001/jamaophthalmol.2015.1468

Brunetti, R., Del Gatto, C., & Delogu, F. (2014). eCorsi: implementation and testing of the Corsi block-tapping task for digital tablets. Front Psychol, 5, 939. 10.3389/fpsyg.2014.00939

Calabrèse, A., To, L., He, Y., Berkholtz, E., Rafianm, P., & Legge, G. E. (2018). Comparing performance on the MNREAD iPad application with the MNREAD acuity chart. In (Vol. 18): Association for Research in Vision and Ophthalmology Inc.

Campbell, F. W., Howell, E. R., & Johnstone, J. R. (1978). A comparison of threshold and suprathreshold appearance of gratings with components in the low and high spatial frequency range. J Physiol, 284, 193–201. 10.1113/jphysiol.1978.sp012535

Campbell, F. W., & Robson, J. G. (1968). Application of fourier analysis to the visibility of gratings. The Journal of Physiology, 197(3), 551–566. 10.1113/jphysiol.1968.sp008574

Chung, S. T. L. (2011). Improving Reading Speed for People with Central Vision Loss through Perceptual Learning. Investigative Ophthalmology & Visual Science, 52(2), 1164–1170. 10.1167/iovs.10-6034

Cowan, N. (2001). The magical number 4 in short-term memory: a reconsideration of mental storage capacity. Behav Brain Sci, 24(1), 87–114; discussion 114-185. 10.1017/s0140525x01003922

Elliot, D. B., Yang, K. C. H., & Whitaker, D. (1995). Visual acuity changes throughout adulthood in normal, healthy eyes: seeing beyond 6/6. In (Vol. 72, pp. 186–191): Optom Vis Sci.

Ferris, F. L., 3rd, Kassoff, A., Bresnick, G. H., & Bailey, I. (1982). New visual acuity charts for clinical research. Am J Ophthalmol, 94(1), 91–96.

Habtamu, E., Bastawrous, A., Bolster, N. M., Tadesse, Z., Callahan, E. K., Gashaw, B., Macleod, D., & Burton, M. J. (2019). Development and Validation of a Smartphone-based Contrast Sensitivity Test. Transl Vis Sci Technol, 8(5), 13. 10.1167/tvst.8.5.13

Hung, S. C., & Seitz, A. R. (2014). Prolonged training at threshold promotes robust retinotopic specificity in perceptual learning. J Neurosci, 34(25), 8423–8431. 10.1523/jneurosci.0745-14.2014

Ioannidis, J. P. A. (2005). Why Most Published Research Findings Are False. PLOS Medicine, 2(8), e124. 10.1371/journal.pmed.0020124

Jayakumar, S., Maniglia, M., Guan, Z., Green, C. S., & Seitz, A. R. (2024). PLFest: A New Platform for Accessible, Reproducible, and Open Perceptual Learning Research. Journal of Cognitive Enhancement, 8(4), 334–345. 10.1007/s41465-024-00299-w

Kannan, L., Lelo de Larrea-Mancera, E. S., Maniglia, M., Vodyanyk, M. M., Gallun, F. J., Jaeggi, S. M., & Seitz, A. R. (2024). Multidimensional relationships between sensory perception and cognitive aging [Perspective]. *Frontiers in Aging Neuroscience*, Volume 16–2024. 10.3389/fnagi.2024.1484494

Kollbaum, P. S., Jansen, M. E., Kollbaum, E. J., & Bullimore, M. A. (2014). Validation of an iPad Test of Letter Contrast Sensitivity. Optometry and Vision Science, 91(3), 291–296. 10.1097/OPX.0000000000000158

Labiris, G., Panagiotopoulou, E. K., Delibasis, K., Duzha, E., Bakirtzis, M., Panagis, C., Boboridis, K., Mokka, A., Balidis, M., Damtsi, C., & Ntonti, P. (2023). Validation of a web-based distance visual acuity test. J Cataract Refract Surg, 49(7), 666–671. 10.1097/j.jcrs.0000000000001176

Levi, D. M., & Li, R. W. (2009). Perceptual learning as a potential treatment for amblyopia: A mini-review. Vision Research, 49(21), 2535–2549. 10.1016/j.visres.2009.02.010

Liang, J., Zhou, Y., Fahle, M., & Liu, Z. (2015). Specificity of motion discrimination learning even with double training and staircase. J Vis, 15(10), 3. 10.1167/15.10.3

Maniglia, M., Pavan, A., Sato, G., Contemori, G., Montemurro, S., Battaglini, L., & Casco, C. (2016). Perceptual learning leads to long lasting visual improvement in patients with central vision loss. Restorative Neurology and Neuroscience, 34(5), 697–720. 10.3233/RNN-150575

Maniglia, M., Soler, V., & Trotter, Y. (2020). Combining fixation and lateral masking training enhances perceptual learning effects in patients with macular degeneration. Journal of Vision, 20(10), 19–19. 10.1167/jov.20.10.19

Matzen, L. E., Benz, Z. O., Dixon, K. R., Posey, J., Kroger, J. K., & Speed, A. E. (2010). Recreating Raven’s: Software for systematically generating large numbers of Raven-like matrix problems with normed properties. Behavior Research Methods, 42(2), 525–541. 10.3758/BRM.42.2.525

Pahor, A., Mester, R. E., Carrillo, A. A., Ghil, E., Reimer, J. F., Jaeggi, S. M., & Seitz, A. R. (2022). UCancellation: A new mobile measure of selective attention and concentration. Behavior Research Methods, 54(5), 2602–2617. 10.3758/s13428-021-01765-5

Pahor, A., Seitz, A. R., & Jaeggi, S. M. (2022). Near transfer to an unrelated N-back task mediates the effect of N-back working memory training on matrix reasoning. Nature Human Behaviour, 6(9), 1243–1256. 10.1038/s41562-022-01384-w

Pahor, A., Stavropoulos, T., Jaeggi, S. M., & Seitz, A. R. (2019). Validation of a matrix reasoning task for mobile devices. Behavior Research Methods, 51(5), 2256–2267. 10.3758/s13428-018-1152-2

Peñaloza, B., Goddin, T. L., Friedman, D. S., Owsley, C., & Kwon, M. (2025). Age-Related Changes in Mesopic Reading Vision Across Adulthood. Invest Ophthalmol Vis Sci, 66(3), 40. 10.1167/iovs.66.3.40

Peñaloza, B., & Kwon, M. (2025). Significant Age-Related Visual Declines but Preserved Binocular Summation of Contrast Sensitivity and Visual Acuity Across Adulthood. Invest Ophthalmol Vis Sci, 66(14), 63. 10.1167/iovs.66.14.63

Polat, U., Schor, C., Tong, J.-L., Zomet, A., Lev, M., Yehezkel, O., Sterkin, A., & Levi, D. M. (2012). Training the brain to overcome the effect of aging on the human eye. Scientific Reports, 2(1), 278. 10.1038/srep00278

Rono, H., Bastawrous, A., Macleod, D., Wanjala, E., Gichuhi, S., & Burton, M. (2019). Peek Community Eye Health - mHealth system to increase access and efficiency of eye health services in Trans Nzoia County, Kenya: study protocol for a cluster randomised controlled trial. Trials, 20(1), 502. 10.1186/s13063-019-3615-x

Sagi, D. (2011). Perceptual learning in Vision Research. Vision Research, 51(13), 1552–1566. 10.1016/j.visres.2010.10.019

Tan, D. T. H., & Fong, A. (2008). Efficacy of neural vision therapy to enhance contrast sensitivity function and visual acuity in low myopia. Journal of Cataract & Refractive Surgery, 34(4). https://journals.lww.com/jcrs/fulltext/2008/04000/efficacy_of_neural_vision_therapy_to_e nhance.23.aspx

Valentine, T., Block, C., Eversole, K., Boxley, L., & Dawson, E. (2020). Wechsler Adult Intelligence Scale-IV (WAIS-IV). In The Wiley Encyclopedia of Personality and Individual Differences (pp. 457-463). 10.1002/9781118970843.ch146

Watson, A. B., & Ahumada, A. J., Jr. (2005). A standard model for foveal detection of spatial contrast. J Vis, 5(9), 717–740. 10.1167/5.9.6

Wolfe, J. M., & Horowitz, T. S. (2017). Five factors that guide attention in visual search. Nature Human Behaviour, 1(3), 0058. 10.1038/s41562-017-0058

Xiao, L. Q., Zhang, J. Y., Wang, R., Klein, S. A., Levi, D. M., & Yu, C. (2008). Complete transfer of perceptual learning across retinal locations enabled by double training. Curr Biol, 18(24), 1922–1926. 10.1016/j.cub.2008.10.030

Zhang, J.-Y., & Yang, Y.-X. (2014). Perceptual learning of motion direction discrimination transfers to an opposite direction with TPE training. Vision Research, 99, 93–98. 10.1016/j.visres.2013.10.011

Zhang, J. Y., & Yu, C. (2016). The transfer of motion direction learning to an opposite direction enabled by double training: A reply to Liang et al. (2015). J Vis, 16(3), 29. 10.1167/16.3.29

